# Predicting protein-peptide interaction sites using distant protein complexes as structural templates

**DOI:** 10.1101/398768

**Authors:** Isak Johansson-Åkhe, Claudio Mirabello, Björn Wallner

## Abstract

Protein-peptide interactions play an important role in major cellular processes, and are associated with several human diseases. To understand and potentially regulate these cellular function and diseases it is important to know the molecular details of the interactions. However, because of peptide flexibility and the transient nature of protein-peptide interactions, peptides are difficult to study experimentally. Thus, computational methods for predicting structural information about protein-peptide interactions are needed. Here we present InterPep, a pipeline for predicting protein-peptide interaction sites. It is a novel pipeline that, given a protein structure and a peptide sequence, utilizes structural template matches, sequence information, random forest machine learning, and hierarchical clustering to predict what region of the protein structure the peptide is most likely to bind. When tested on its ability to predict binding sites, InterPep successfully pinpointed 255 of 502 (50.7%) binding sites in experimentally determined structures at rank 1 and 348 of 502 (69.3%) among the top five predictions using only structures with no significant sequence similarity as templates. InterPep is a powerful tool for identifying peptide-binding sites; with a precision of 80% at a recall of 20% it should be an excellent starting point for docking protocols or experiments investigating peptide interactions. The source code for InterPred is available at http://wallnerlab.org/InterPep/

## Introduction

Protein-protein interactions are a fundamental part of all major biological processes. A particularly interesting class of protein-protein interactions are those involving interaction including intrinsically disordered regions. These regions are often the size of small peptide fragments 5 to 25 residues long^1^ and part of proteins involved in regulation, recognition, and signaling requiring dynamic and specific responses^2, 3^. When investigating these interactions, it is common practice to isolate the binding motif of the disordered region and analyze the binding as a protein-peptide interaction^4, 5^. Protein interactions involving disordered regions, as well as other protein-peptide interactions, are frequently associated with various human diseases such as cancer, cardiovascular disease, amyloidosis, and neurodegenerative diseases^6, 7^. It has also been shown that protein-peptide interactions can be regulated using small molecules^8–10^, making them prime candidates as drug targets. Thus, understanding the structural details of protein-peptide interactions is fundamental for our understanding of complex cellular processes and complex diseases, and in addition they can provide a basis for rational drug design to modify protein-peptide interactions.

Experimental methods for determining protein-protein interactions in molecular detail, such as x-ray crystallography, cryo-EM, and NMR, are both difficult and time-consuming^11^. High-throughput methods that measure binary interaction, such as affinity-purification mass spectrometry, yeast-two-hybrid, and BioID^12–14^ are lacking in molecular detail. Thus, many computational methods have been developed to predict protein complex interactions^15^. There are both methods that predict binary interactions from the amino acid sequence^16^ or by combining other sources of information^17^, as well as methods that predict the complete 3D-structure of a complex using sequence- and profile-based alignments^18, 19^ or structural alignments^20, 21^ to libraries of interacting proteins of known structure.

Methods that utilize structural alignments have proved to be especially successful in the latest CASP12-CAPRI^22^. These methods search through the Protein Data Bank^23^ for interacting complexes structurally similar to both query proteins, and work well when both partner proteins are well-ordered protein structures. However, for modeling protein-peptide interactions these methods cannot be used, since the interaction in this case is between an ordered partner and a smaller, relatively flexible, peptide fragment. Instead, methods that try to dock the peptide onto the ordered partner have been applied to this problem with some success for very short peptides (*≤*4 residues)^24^. The success for larger peptides is limited due to the large number of peptide conformations and binding poses that needs to be sampled^25–27^.

The sampling problem can remedied to some degree by coarse-graining using side-chain centroids like in Rosetta FlexPep-Dock^25^ or the CABS force field in CABS-dock^27^. Another way to tackle the sampling problem is to first predict the peptide binding site on the protein surface^28–32^. If successful, docking methods can focus the attention to the predicted binding site instead of wasting time on sampling incorrect binding sites. In addition, experiments can be directed towards the predicted binding site and the functional effects of the interaction can be studied. PepSite^28, 29^ is a method that predicts the peptide binding sites by describing he preferred binding environment for peptide residues in protein-peptide interactions with known structure using a spatial position-specific scoring matrix (S-PSSM) that can be used to scan the protein surface for candidate binding sites for peptide residues. More recent methods, PeptiMap^30^ predicts peptide binding by adapting the fragment mapping (FTmap) for ligand peptide binding site prediction^33^ to peptide binding characteristics, PEP-SiteFinder^31^ combines peptide 3D conformation generation and fast rigid body docking, and ACCLUSTER^32^ clusters surface residues that have good chemical interactions with amino acid probes.

Other methods combine both the peptide binding site prediction with peptide-protein docking into one pipeline. GalaxyPepDock^34^ performs template-based docking using templates from a database of experimentally determined peptide-protein structures and builds models using molecular docking, while PIPER-FlexPepDock^35^ combines coarse-grained mass-sampling of the protein receptor surface using PIPER^36^ with molecular docking of the peptide using Rosetta FlexPepDock^25^.

PeptiMap^30^, PEP-SiteFinder^31^, and ACCLUSTER^32^ are unfortunately available only as web servers with limited program-matical access. Similarly, PIPER-FlexPepDock^35^ is only available either through a web-server or an expensive license. In fact, to the best of our knowledge the only software that is currently available as stand alone download is GalaxyPepDock^34^, and even this method is distributed with precompiled executables, meaning it only runs on a select few system architectures. The lack of method availibility clearly hinders progress in the field and makes it difficult to benchmark against these methods.

In this study, we present InterPep, a novel protein-peptide interaction site prediction pipeline that predicts which residues on the surface of a target protein are most likely to interact with a given peptide fragment. It is based on the hypothesis that, if a protein-peptide interaction exists within the PDB describing a similar interaction surface, that interaction surface can be used as a template to model the interaction. Apart from the use of machine learning another difference to previous studies is that we expand the possible templates for interaction from protein-peptide to any protein-protein interaction. This is a reasonable assumption since protein-peptide interactions adopt the same structural motifs as monomeric protein folds upon binding^37^. Recent studies have suggest that the protein-protein interaction space in current PDB is close to complete^38, 39^, indicating that it could potentially be possible to model the majority of protein-peptide interactions using structural templates.

The complete InterPep pipeline runs mass structural alignments to find interaction templates, ranks these templates with a random forest classifier using both structural and sequence information, and outputs both a predicted per residue probability and an amino acid preference profile for the interaction. Both these types of information could potentially be used as constraints in local protein-peptide docking, such as by Rosetta FlexPepDock^25^.

The performance of InterPep demonstrates the strength of a structural template-based approach to investigate protein-peptide interactions. Even though InterPep certainly can be used to model ordered interactions, we believe that best use-cases will be modeling of interactions with disordered regions, i.e an ordered protein interacting with a disordered partner. These interactions are immensely important but yet extremely difficult to study. The approach can easily be modified and extended to include other types of ligand binding, such as binding of phosphorylated targets and saccharides. InterPep is developed using only freely available software, and the source code is available for download, see link above.

## Methods

### Dataset

To develop and evaluate InterPep, a dataset of known protein-peptide interactions was constructed using Protein Data Bank^23^ (May 19, 2016). The dataset consisted of structures containing interactions between one larger protein chain and a smaller poly-peptide chain of 5 to 25 residues acquired through NMR or X-ray crystallography with a resolution of at least 3Å. Additionally, the shared surface between the peptide and the protein had to be at least 400Å^2^. The peptide length and shared surface area requirement were based on the MoRF definitions from Mohan *et al.* (2006). Peptides of these lengths have been observed to be disordered when unbound, representing intrinsically disordered and difficult-to-predict peptides or regions. The size of the shared surface was calculated with NACCESS^40^. Furthermore, to focus on surface interaction, any cases where the peptide was buried by more than 60% were filtered. To ensure common proteins were not overrepresented within the dataset, the larger protein chains were clustered at 30% sequence identity, using BLASTCLUST^41^. The final dataset consisted of 502 protein-peptide interactions, and is freely available at http://wallnerlab.org/InterPep.

Several of the protein-peptide complexes in the dataset are taken from multi-chain structures containing addititional protein chains. This was a deliberate choice, since when investigating protein-peptide interactions it is often not known if the receptor protein oligomerize or interacts with additional proteins. As such, it is important for a protein-peptide interaction site predictor not to falsely predict protein-protein interaction sites. When testing InterPep and other methods, no knowledge of any protein-protein interaction sites was utilized in order to test their capability to predict the correct site, in the presence of potentially disturbing noise from other protein-protein interaction sites.

### Interaction Template Library

An interaction template library was constructed using all pairs of interacting chains from the PDB (May 19, 2016), defined as two chains with any heavy atom within 6Å. Templates where the receptor was less than 25 residues large were removed, as was templates from files with excessive number of chains, such as complete virus capsid structures. For NMR structures, only the first model was included. The interaction template library consisted of 402,102 template pairs of interacting protein chains.

### Protein 3D-models for training and benchmarking

To investigate the impact of model quality of the target protein and to determine whether InterPep is robust enough to work on modeled structures rather than on native structures, a set of structural models derived from the protein sequences of the protein-peptide interaction dataset set was constructed. These models were produced using HHblits^42^ with E-value cutoff at 10^-2^ and maximum pairwise identity of 90%, to search against the `uniprot20_2016_02` database to build the Hidden Markov Model profiles, which were then used to search against the PDB^23^ database clustered at 70% sequence identity. To ensure having models of varying quality, up to eight alignments per target were chosen and modeled by MODELLER^43^. In total, 1,617 models for all targets were constructed in this way.

### InterPep construction

The complete InterPep pipeline is outlined in Figure 1. It consists of four distinct steps: 1. Mass Structural Alignment, 2. Feature Extraction, 3. Random Forest Classification, and 4.Clustering, described in detail below.

**Figure 1.**
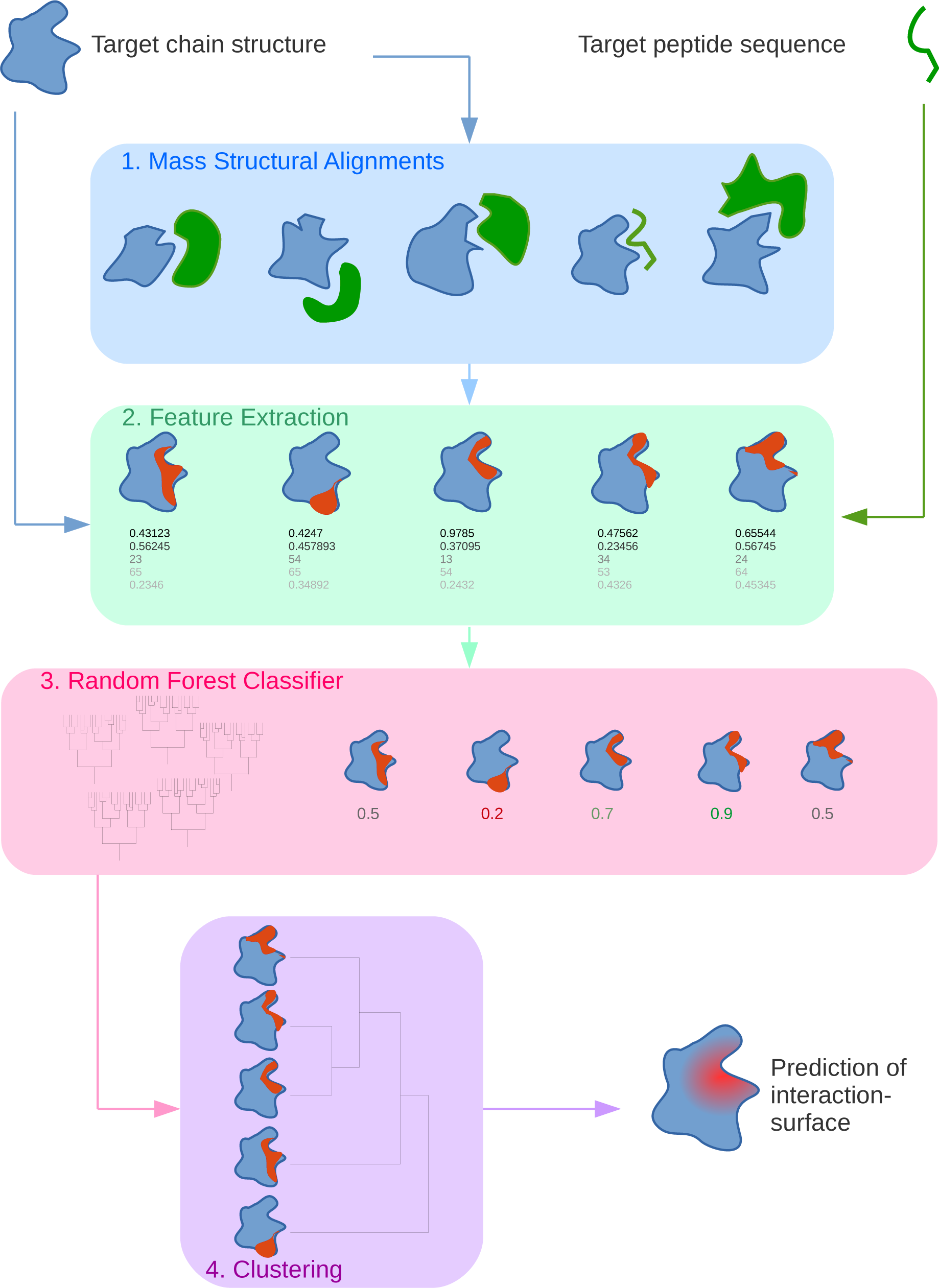
Overview of the InterPep pipeline.

#### 1. Mass Structural Alignment

In the first step, the target protein is structurally aligned using TM-Align^44^ to every protein chain in each pair from the *Interaction Template Library* described above. Any alignment with a TM-score *≥*0.5, which is the cutoff value for a significant structural match^45^, is used as a potential template for the target protein-peptide interaction. The alignment of the residues from the template involved in protein-protein interaction to the equivalent residues in the target protein, gives a template for a potential interaction surface in the target.

#### 2. Feature Extraction

In the second step, features relevant to protein-peptide interactions are calculated for each template. The features used by InterPep are summarized in Table 1, and are described in detail in the Supplementary Information.

**Table 1.** Summary of the features used by InterPep.

| Feature Name | Number of parameters |
| --- | --- |
| **Length** |  |
| Target Peptide | 1 |
| Target Protein Chain | 1 |
| Aligned Template Chain | 1 |
| Aligned Segment | 1 |
| Aligned Interacting Residues | 1 |
| **Alignment Quality** |  |
| Target TM-Score | 1 |
| Template TM-Score | 1 |
| RMSD | 1 |
| **Aligned Region Complexity** |  |
| Contact Order | 1 |
| Longest aligned α-helix | 1 |
| **Amino Acid Composition Distance** |  |
| Between Target Peptide and Template Peptide | 1 |
| Between Target Protein and Template Protein | 1 |
| **Secondary Structure** |  |
| Target Peptide (predicted) | 3 |
| Peptide Template Surface | 3 |
| **Surface Information** |  |
| Relative Exposure of Interaction Surface | 1 |
| Conservation of Interaction Surface | 1 |
| **Template Peptide Information** |  |
| Mean Sequential Distance | 1 |
| Median Sequential Distance | 1 |
| Summed representative lengths | 1 |
| **Model Information (optional)** |  |
| Model Quality measured by Sequence Identity of Alignment | 1 |

#### 3. Random Forest Classification

In the third step, the extracted features are used in a random forest classifier to separate correct and incorrect templates. Templates were considered correct if at least 80% of the interaction site residues suggested by the template where correct. Other cutoffs where also tried, including training regressional random forest to predict the fraction of correct residues, but classification was superior (see Supplementary Information). The random forest is implemented using scikit-learn^46^ with 100 trees and no depth limit. To avoid overfitting, a maximum of 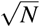 features were considered in each branching, where *N* is the total number of features. The random forest was trained using jack-knifing to estimate the performance. The jack-knifing was performed such that when testing on one protein-peptide pair, templates from all other targets, except those which matched with a TM-score*≥*0.5 to the test protein, were used for training. In addition, to avoid biasing the data toward common structural motifs, when training the Random Forest every protein-peptide pair could only contribute up to 2,000 randomly selected templates to the training data. The output of the random forest for a given template is the probability that the template is correct, given by the definition above.

#### 4. Clustering

In the fourth and final step, the potential interaction surfaces are clustered using hierarchical clustering utilizing the SciPy package^47^, and the average method for calculating distances to clusters. The distance between two potential interaction surfaces is calculated as the mean of the distances between the coordinates for each residue in the different surfaces. The distance cutoff is determined automatically through the elbow method^48^. The highest predicted probability among the templates in a cluster defines the score of that cluster (*cluster score*).

The top 10% of all templates within the highest scoring cluster are then combined through a weighted average to produce the individual local residue scores that make up the final prediction. The weights are the probabilities predicted by the random forest for the templates, and *local score* measures the likelihood of each residue to be involved in a protein-peptide interaction. Hierarchical clustering and the 10% cutoff were utilized in favor of calculating a weighted average over all structures to ensure that strong and correct signals were not hidden by many weak ones, and to make the final prediction more robust to random outliers and fragmented templates by basing it on an ensemble of predictions.

#### InterPep score

For every prediction method, it is important to have a score that correlates with the confidence in the predictions. In InterPep, the *global score* is designed to measure the confidence in the prediction, and is defined by combining the *cluster score* with the highest *local score* using a weighted mean, selecting the weights 0.61 for *cluster score* and 0.39 for *local score* maximized the correlation to 0.63 for the quality of prediction as measured by Matthew’s Correlation Coefficient (MCC) between correct and incorrect template residues, Figure 2a. The *global score* has the advantage that it combines the confidence in the predicted area measured by the *cluster score* with the confidence of the predictions in that area measured by the *local score*, and should thus represent the confidence in the prediction as a whole. As *cluster score* indirectly describes the maximum quality of the templates available, it is natural that it also has the highest weight. The distribution of precisions (fraction of correct residues) for different score thresholds shows that the *global score* is a good estimate of performance, see Figure 2b. For *global score* above 0.4 the majority of the predictions have precisions above 60% increasing to 80%, with very few predictions with precision less than 50% for *global score* above 0.7.

**Figure 2.**
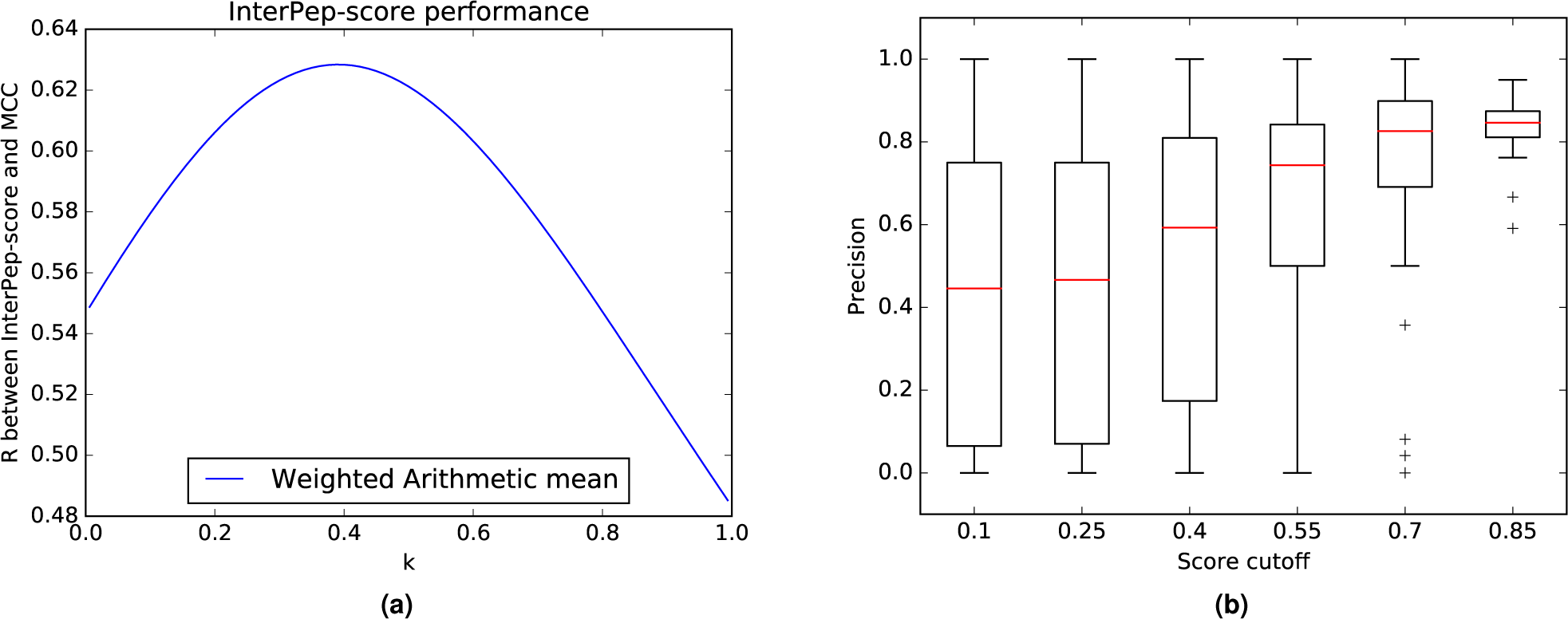
(a) Correlation between InterPep *global score* and prediction quality measured as Matthew’s Correlation Coefficient between the local score and native contacts, for different ways of combining the *cluster score* and highest *local score*. The weight for the *local score* is *k* and *cluster score* 1 *− k*. Optimal weight is found at *k* = 0.39. (b) Distribution of precision of predictions for different InterPep score cutoff values.

## Results and Discussion

### Development of InterPep

The overall method has been described in Methods. Below we describe, benchmark, and validate some of the design choices that were made during the development of InterPep.

#### Interaction Template Library Analysis

InterPep relies on finding interaction templates that can be used to model a particular protein-peptide interaction. As such, the set of interaction templates, or the interaction template library, available to InterPep sets the boundaries for the prediction. Relevant questions in this regard are how many protein-peptide interactions can be modelled by such templates, and if protein-protein interaction templates can be used to model protein-peptide interactions? To answer these questions, the full interaction template library (see Methods) was divided into a peptide-binding and a non-peptide-binding part, and the possibility of using perfect combinations of templates from the full, peptide and non-peptide parts to model protein-peptide interaction was assessed. Perfect combinations of templates for the different template libraries were constructed by fitting all templates available to the correct answer using linear regression, as shown in the equation below:

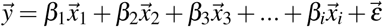

Where 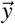 is a vector representation of the target protein containing binary values denoting which residues are involved in protein-peptide interaction in the native structure and which are not, *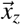* is the interaction vector of template *z*, *β*_*z*_ is a constant, *i* is the total number of templates, and *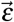* is a vector of errors. All *β* are assigned values so that the total error 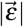 is as small as possible. 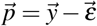 gives the best possible prediction available, and is a prediction vector similar to the *local score* from InterPep.

A binding site was considered correctly identified if it had at least 50% overlap with native interaction site. All templates with BLAST E*<*0.001 to the targets had been removed from the Interaction Template Library prior to this analysis. As can be seen from Table 2, there are templates available to model the majority of the targets (81.3%) from the test set using all interactions in the template library. By restricting the library to only non-peptide-binding templates, it is still possible to model 67.3% of the targets. This increases to 73.5% when instead using only peptide-binding templates. However, the fact that it is possible to model a substantial proportion of the peptide interactions with the non-peptide interaction templates and that more targets can be modeled using all templates (both peptide and non-peptide) supports the hypothesis that protein-protein interfaces can be used to predict protein-peptide interfaces.

**Table 2.** Number of true sites which can be described as combinations of templates. Number of peptide interactions found in the test set containing 502 interaction sites, depending on the used template library. *All templates* correspond to all templates in the interaction library, *Only peptide-binding templates* is the subset of all templates that involve peptide interactions, *Only non-peptide templates* is the subset of all templates that do not involve peptides,*Random* is what would be expected by chance, and was generated by randomly predicting *n* number of residues on the surface of the protein as interacting, where *n* is the number of interacting residues in the native structure, *Total* is the total number of sites in the test set. No templates matching the target with BLAST E*<*0.001 were used.

| Template library | # protein-peptide interactions that can be modeled |
| --- | --- |
| All templates | 408 (81.3%) |
| Only peptide-binding templates | 369 (73.5%) |
| Only non-peptide templates | 340 (67.7%) |
| Random | 54 (10.8%) $\sigma=5.85$ |
| Total sites | 502 (100.0%) |

### Comparison to Established Methods

The performance of InterPep was benchmarked against other established methods. There are several published methods that potentially could be included in a benchmark. However, as most of them are only available as web servers with limited or even blocked programmatical access, we only included the following methods that could actually run on the full benchmark set: PepSite2^29^, PeptiMap^30^, and GalaxyPepDock^34^, of these the latter is available as stand alone download, while the other two allows programmatical web access. In addition, two reference methods were also included, one method using surface conservation (Surf-Cons) calculated using Rate4Site^49^ for surface residues, and one method similar to InterPep, but with the random forest classifier score replaced with the TM-score from TM-align (InterPep-TMonly).

Since some methods give very few predictions and some very similar predictions for the different targets we restricted the analysis to the first ranked prediction for each target. For each method, the top-1 prediction for each targets of the dataset was analysed in Precision-Recall curve, see Figure 3, where the overall precision of the method is plotted against what fraction of true hits that are found for varying thresholds. Unfortunately, even though PeptiMap calculates several scores that could potentially be used as threshold cutoffs^30^, none of these scores are available to the user and thus PeptiMap could only be represented by a single point. PepSite2 only allows peptides 10 residues long or less, to make sure PepSite2 was not disadvantaged for longer peptides, it was run with a sliding window of size 10 for the whole peptide and the best prediction was chosen for each target. Also, Surf-Cons can sometimes predict several different binding sites on the same target, in these cases only the best was included.

**Figure 3.**
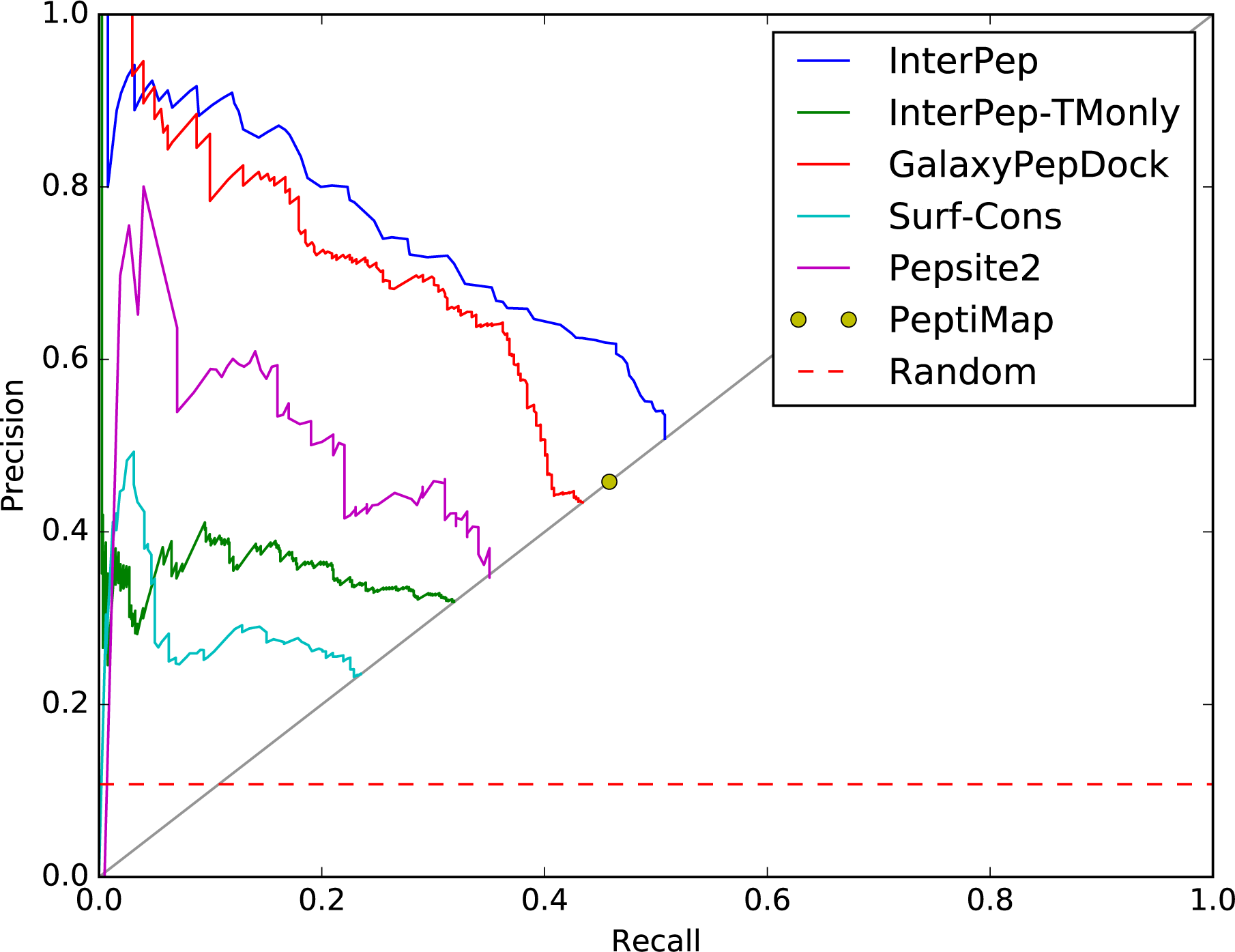
Precision-recall measuring the ability of different methods to correctly identify the correct binding sites at top-1. Precision is the fraction of predictions so far made which have been correct. Recall is the fraction of targets for which the correct site has been identified. The curves stop when predictions are made for all targets at the line Precision=Recall, since only one prediction per target is considered.

Overall, InterPep performed best of all methods considered, with a consistently larger precision over 5% recall compared to all other methods, see Figure 3. For 20% recall, InterPep has a precision of 80%, compared to 73% for GalaxyPepDock, 61% for Pepsite2, 36% for InterPep-TMonly, and 26% for Surf-Cons. InterPep also has the highest maximum recall of all methods: 50.7% corresponding to 255 out of 502 targets, tabulated in Table 3 for easier reference. Note that PeptiMap, which did not provide a score to be used for a curve in Figure 3 has a slightly higher maximum recall compared to GalxyPepDock, 45.8% vs. 43.4% corresponding to 230 and 218 correct predictions, respectively.

**Table 3.** Number of correct sites identified. The number of target pairs where the correct peptide-binding site was identified in the top scorer (cluster for InterPep) for the different methods. *#protein-peptide interaction sites* measures the number of predictions that hit a protein-peptide interaction site, while *#protein-protein interaction sites* measures the number of predictions that hit a protein-protein interaction site that is an interaction but not the correct protein-peptide interaction site. *Random* is what would be expected by chance, and was generated by randomly predicting *n* number of residues on the surface of the protein as interacting, where *n* is the number of interacting residues in the native structure. *Total* is the total number of sites in the test set. ^*\**^For template methods, no templates matching the target with BLAST E*<*0.001 were used. ^*\*\**^To make sure PepSite2 was not disadvantaged for longer peptides, it was run with a sliding window of size 10 for the whole peptide and the best prediction was chosen.

| Method | # protein-peptide interaction sites (maximum recall) | # protein-protein interaction sites |
| --- | --- | --- |
| InterPep* | **255 (50.7%)** | 32 (14.2%) |
| PeptiMap | 230 (45.8%) | 39 (17.3%) |
| GalaxyPepDock* | 218 (43.4%) | **24 (10.6%)** |
| PepSite2** | 176 (35.0%) | 54 (23.9%) |
| InterPep-TMonly* | 160 (31.9%) | 48 (21.2%) |
| Surf-Cons | 118 (23.5%) | 32 (14.2%) |
| Random | 54 (10.8%) $\sigma=5.9$ | 20 (8.8%) $\sigma=2.6$ |
| Total sites | 502 (100.0%) | 226 (100.0%) |

Table 3 also reports the number of times a protein-protein interaction site is incorrectly predicted to be a protein-peptide site. All methods except GalaxyPepDock have a significantly higher number of these types of mistakes than would be expected by chance (26 is the limit for p=0.05). GalaxyPepDock is using a template database from PeptiDB^50^ that only contains protein-peptide interactions, thus it should be less likely to find protein-protein interaction sites than the other template-based methods InterPep and InterPep-TMonly, which also takes protein-protein interactions as templates. However, this inclusion of protein-protein interaction-templates significantly improved InterPep’s recall when locating protein-peptide interaction-sites; 60 of the 255 interfaces correctly identified by InterPep used a protein-protein interaction-site as the highest ranked template, which puts InterPep’s recall above GalaxyPepDock’s, which identified only 218 sites.

Furthermore, as protein-peptide interactions are similar to protein-protein interactions^37^, it makes sense that most other methods should produce a higher than random amount of errors mistaking protein-protein interfaces for protein-peptide interfaces. Pepsite2 reports the highest number of these errors, which is unsurprising, as its site-prediction protocol is based on finding “hot-spots” for binding different amino-acids by using position-specific scoring matrices of the amino acids’ preferred surroundings, and then finding a site where these hot-spots chain together. As the environments of amino acids in protein-peptide interactions are similar to those in protein-protein interactions^37^, it makes sense that a method based around finding such environments would struggle to differentiate protein-protein from protein-peptide interaction-sites.

The fact that InterPep performs much better than InterPep-TMonly clearly indicates that the random forest classifier in InterPep is much better at ranking predictions compared to TMscore alone. Still, even tough InterPep-TMonly does a poor job in ranking predictions between targets, it achieves a decent maximum recall of 31.9% (160/502 correct predictions) comparable to PepSite2 with 35% (176/502 correct predictions). The reason why TM-score is not a good discriminator by itself is of course that a high TM-score, ergo a close structural match, is important, but is not alone sufficient, to classify a prediction as correct, and additional information about the peptide and sequences are needed to make the prediction more accurate. This also explains why InterPep-TMonly makes mistakes by finding more protein-protein interaction sites compared to InterPep (48 vs. 32), as without additional information about the template and peptide it is impossible to tell the difference between protein-protein and protein-peptide interaction templates using only TM-score, see Table 3.

### InterPep on Model Structures

Thus far, InterPep has only been trained and benchmarked on native structures. Of course, in a real case scenario it is more than likely the prediction needs to be performed using modeled structures, simply because the native structure does not exist for the protein target of interest. It is therefore important to benchmark and assess the performance also on modeled structures.

Models of the native structures were constructed (see Methods) and used as input to the InterPep pipeline. The ability of InterPep to predict the correct binding sites of model structures was only slightly lower than for native structures, correctly identifying 49.8% protein-peptide interaction sites in models compared to 50.7% for native structures, see Table 4. Since InterPep was only trained on native structures it seems to suffer a minor performance loss when dealing with model structures. To circumvent this problem, InterPep was retrained using both native and modelled structures (InterPepM). InterPepM also includes the sequence identity between the target and the model template as an additional feature of model quality, for the random forest classification step. The retrained version performs slightly better for models (52.2% vs. 49.8%), and slightly worse on native structures (48.4% vs. 50.7%). In addition, the InterPepM score correlates slightly better with performance (MCC) for the model structures than the original InterPep trained only for native structures (R=0.530 vs. R=0.523, see Supplementary Information). This indicates that there is a difference in how the random forest evaluates templates depending on whether they originate from alignments to modelled or native structures.

**Table 4.** Number of correct protein-peptide interaction sites identified. The number of target pairs where the correct peptide-binding site was identified in the top cluster, when trained on native structures or on native and model structures, and tested on native and model structures, respectively.

|  | Tested on natives | Tested on models |
| --- | --- | --- |
| InterPep | 255 (50.7%) | 805 (49.8%) |
| InterPepM | 243 (48.4%) | 844 (52.2%) |
| Total sites | 502 (100.0%) | 1617 (100.0%) |

Finally, we investigated whether InterPepM performance on native and modelled structures was related to the quality of the models, e.g. whether InterPep performs well for the native structure, and whether poor performance could be attributed to poor model quality. We analyzed the difference in precision between the models and their corresponding native structure as a function of the quality of the modelled structures measured by S-score^51^, see Figure 4. We found that there was very small correlation between model quality and the difference in performance between model and native (R=0.10), and that the difference in performance was often minor (mean=-0.14). This is good news, since it means that the predictions are robust to modelling errors. However, it is also apparent that when comparing the global InterPep score of the natives to their respective models, the predictions for models that have worse InterPep score than the native also have a significantly (P<1.1E-5) larger performance difference (mean=-0.26) compared to models with InterPep score better than native (mean=-0.12), lending further credibility to the global InterPep score.

**Figure 4.**
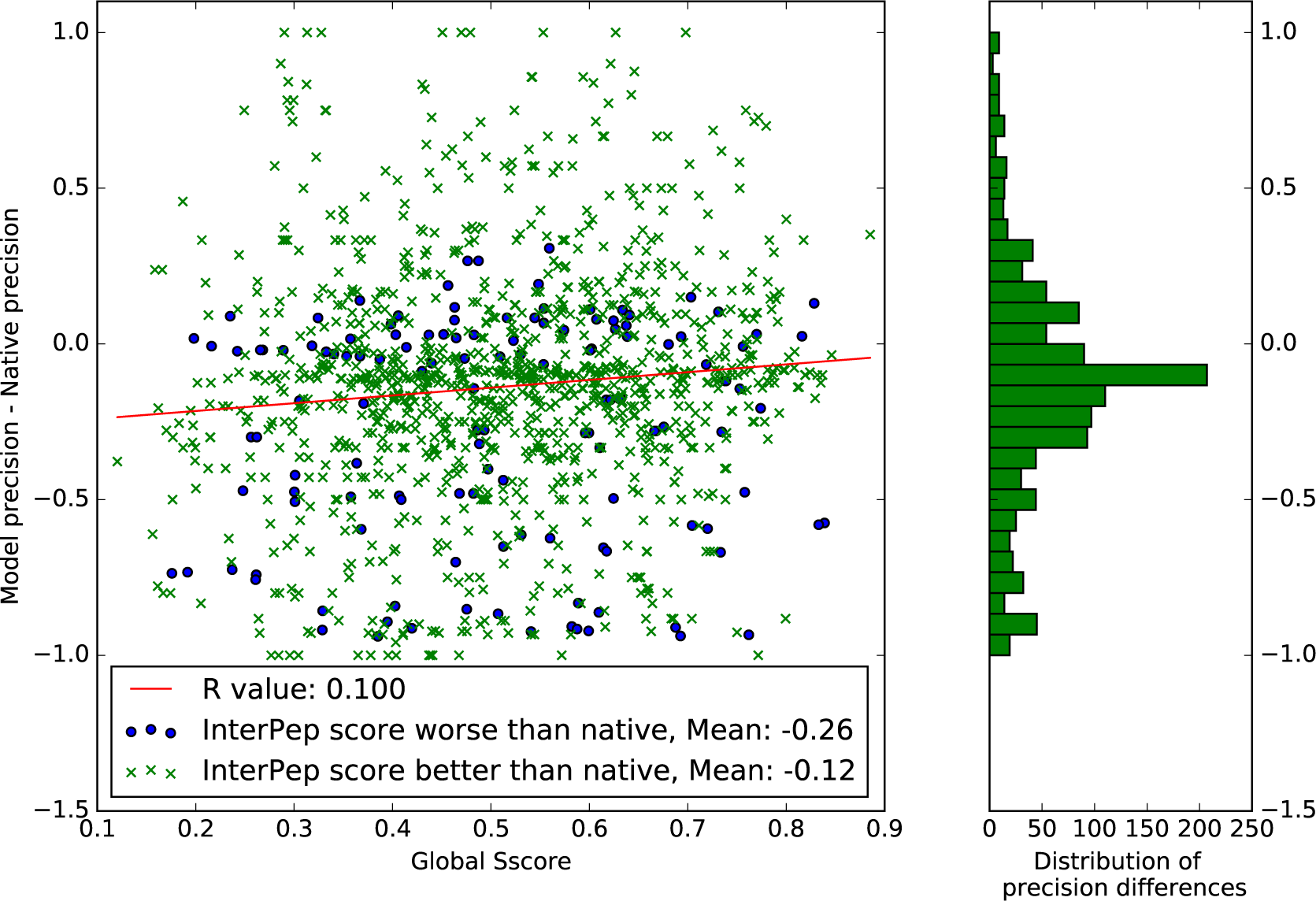
Model quality versus difference in InterPepM precision for model and native. The differences in precision have a mean of −0.138, and a standard deviation of 0.38. The uninteresting cases where predictions for both native and model structures produced a 0 precision result were not included.

Since neither InterPep nor InterPepM is universally better, the InterPep pipeline is distributed with both versions, and depending on what input data is available the appropriate version can be selected.

### Feature Importance Analysis

To get an idea of the degree to which the different features influence the prediction, the relative importance of each feature used by the random forest classifier in InterPep was estimated by Gini impurity (see Supplementary Information for details on how this is calculated). These relative feature importances are shown in Figure 5.

**Figure 5.**
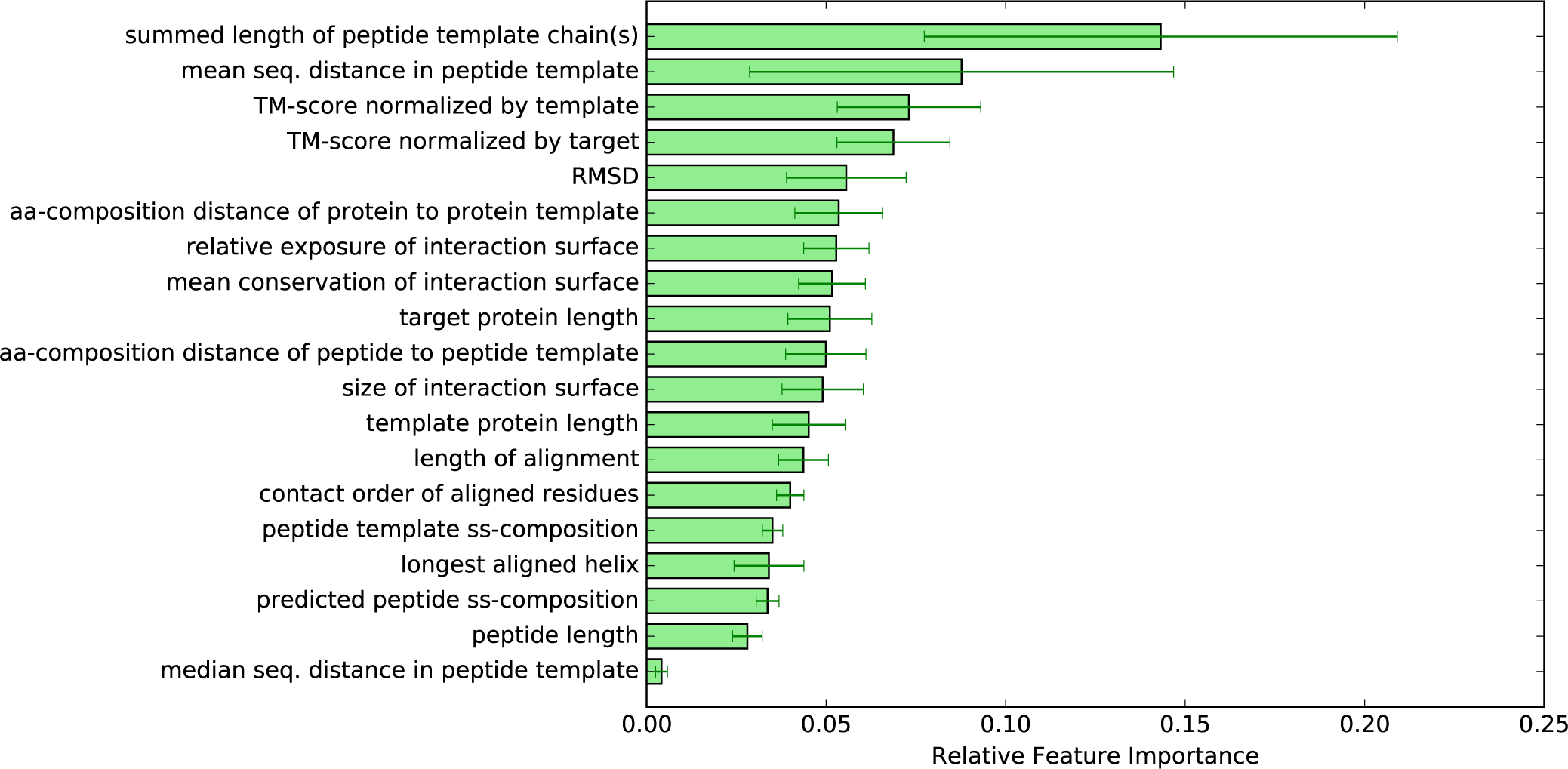
Bar graphs showing the relative importances of the features, calculated by reduction of gini impurity at relevant branchings. These are the mean importances calculated between all trees of the forest.

Of the five most important features, two describe length and mean sequential distance of the peptide template, and the other three describe the quality of the structural alignment between the target and template, see Figure 5. Since the first two features describe how long and how continuous the peptide template is, they can be used to distinguish between whether the interaction partner in the template is a smaller peptide fragment or a longer protein. Indeed, analyzing the splits of these features indicates that a more peptide-like interaction partner for the template means the random forest will generally grant a higher score: 64% of all branchings on peptide template length favored smaller templates, and 56% of branchings on mean sequential distance in the peptide template also favored lower values. This, together with the importance of these features, indicates that peptide-binding templates are more favorable than protein-binding templates when modeling protein-peptide interactions.

Additionally, since structure has been shown to be more conserved than sequence^52^, and is intrinsically linked to function^53^, it is no surprise that features regarding how well the target structurally matches a template should be ranked as important; 56% of TM-score branchings prefer higher TM-score, and 55% of RMSD branchings prefer low RMSDs.

It should be noted that counting label distribution at branchings only allows a very rough and generalized estimate of how a forest estimates a feature, as the total prediction trees are much more complex, e.g. in how the different branchings affect others further down the trees.

### Prediction Examples

Three examples of final predictions from InterPep are shown in Figure 6, to further highlight the strengths and weaknesses of the method. The first example illustrates a high-scoring prediction (InterPep=0.84) in Figure 6a, the structure is colored from red to blue based on the local InterPep score, where red represents regions predicted to be peptide interacting. This is a highly accurate prediction with both high precision and recall of 0.92 and 0.85 respectively, and the predicted peptide-binding region closely resembles the correct peptide-binding region from the native structure shown in red in Figure 6b. It only failed to predict the interaction of the loop close to the N-terminal of the peptide and incorrectly predicted a loop interaction near the C-terminal of the peptide. The next example illustrates another high, but not extreme, InterPep score (InterPep=0.73), prediction in Figure 6c. In this case the interaction surface is correctly identified, i.e. it has very a high precision of 1.00, but is only partly predicted, i.e. it has a low recall of 0.36, compared to the much larger correct surface illustrated in the native complex Figure 6d. The last example illustrates lower scoring InterPep predictions (InterPep=0.46-0.42) and is shown in Figure 6e and g. In this case, the target protein is a homo dimer, and the highest ranked InterPep prediction is actually the dimer interface rather than the true protein-peptide-binding site, see Figures 6e and 6f. The top prediction incorrectly predicts the dimer interface to be the peptide interface. If we know the details of the dimer interface of this protein, and know or suspect the peptide is not competing with it, it would be natural to explore other predictions based on lower ranked clusters when analyzing the results. As can be seen in Figure 6g, the second to highest ranked interface is in the same area as the correct peptide-binding interface, even though the prediction is slightly off the mark, which results in low precision and recall of 0.33 and 0.10, respectively. This example highlights that when using InterPep it is wise to evaluate more than only the highest scoring prediction, especially if the lower scoring predictions have similar scores. In general, it is possible to improve the detection of correct sites from 50.7% for first ranked to 69.3% for the top five ranked predictions, see Figure 7.

**Figure 6.**
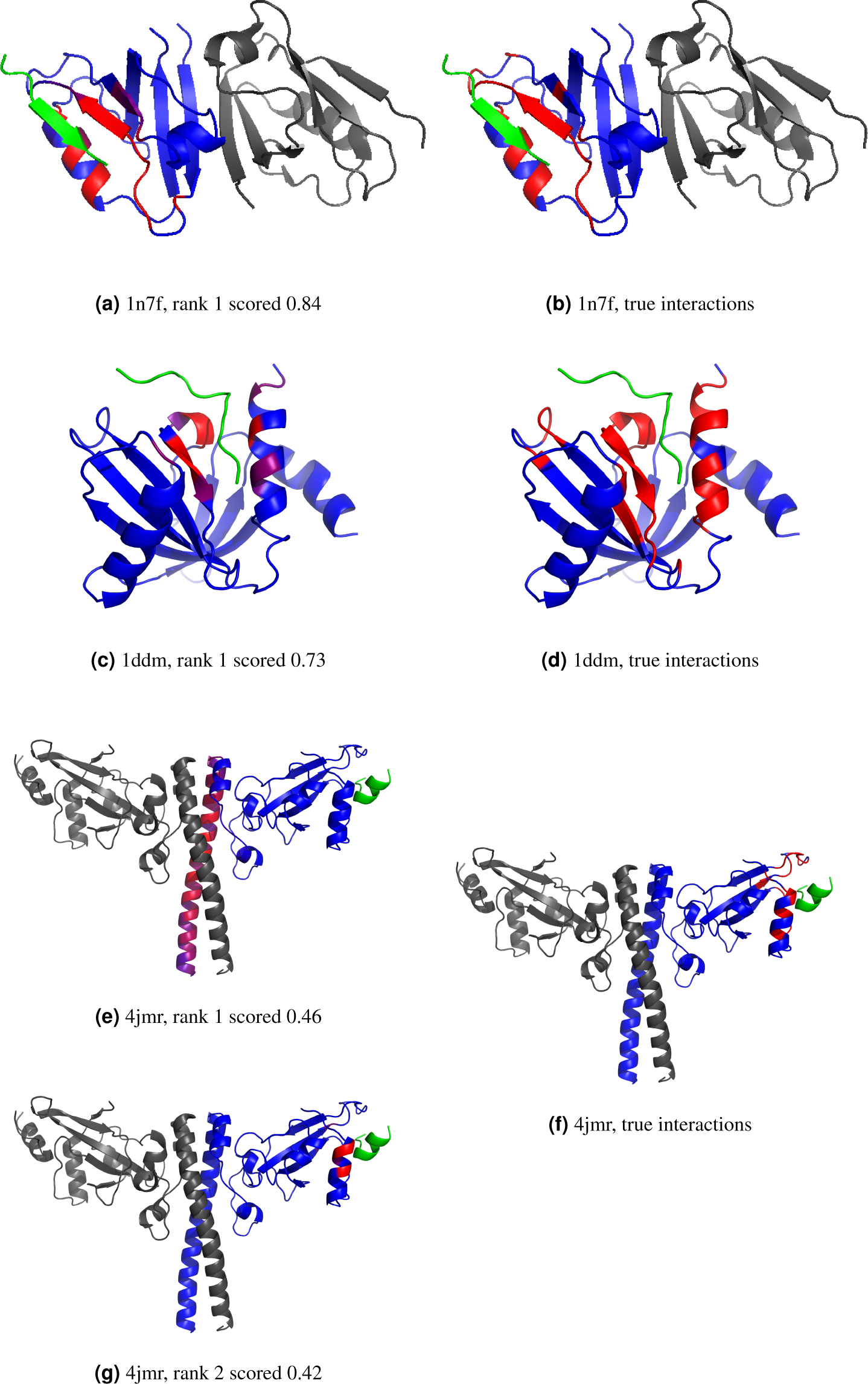
Examples of InterPep predictions shown in native structures. The target protein chain is blue, peptides interacting with the targets are green, and other chains present in the native structure are gray. No information on these additional chains was given during the prediction process. InterPep predictions (a, c, e, and g); the residues of the target protein chain are colored by InterPep prediction values, with red representing strong confidence in interaction, and blue representing no confidence in interaction. The correct peptide interactions (b, d, and f); the residues of the target protein chain which interact with a peptide are marked in red the rest is in blue. Note that all images are of the native structure, and only colored by the prediction from InterPep. InterPep does not predict the structure of complexes, but rather which residues are likely to be involved in interaction.

**Figure 7.**
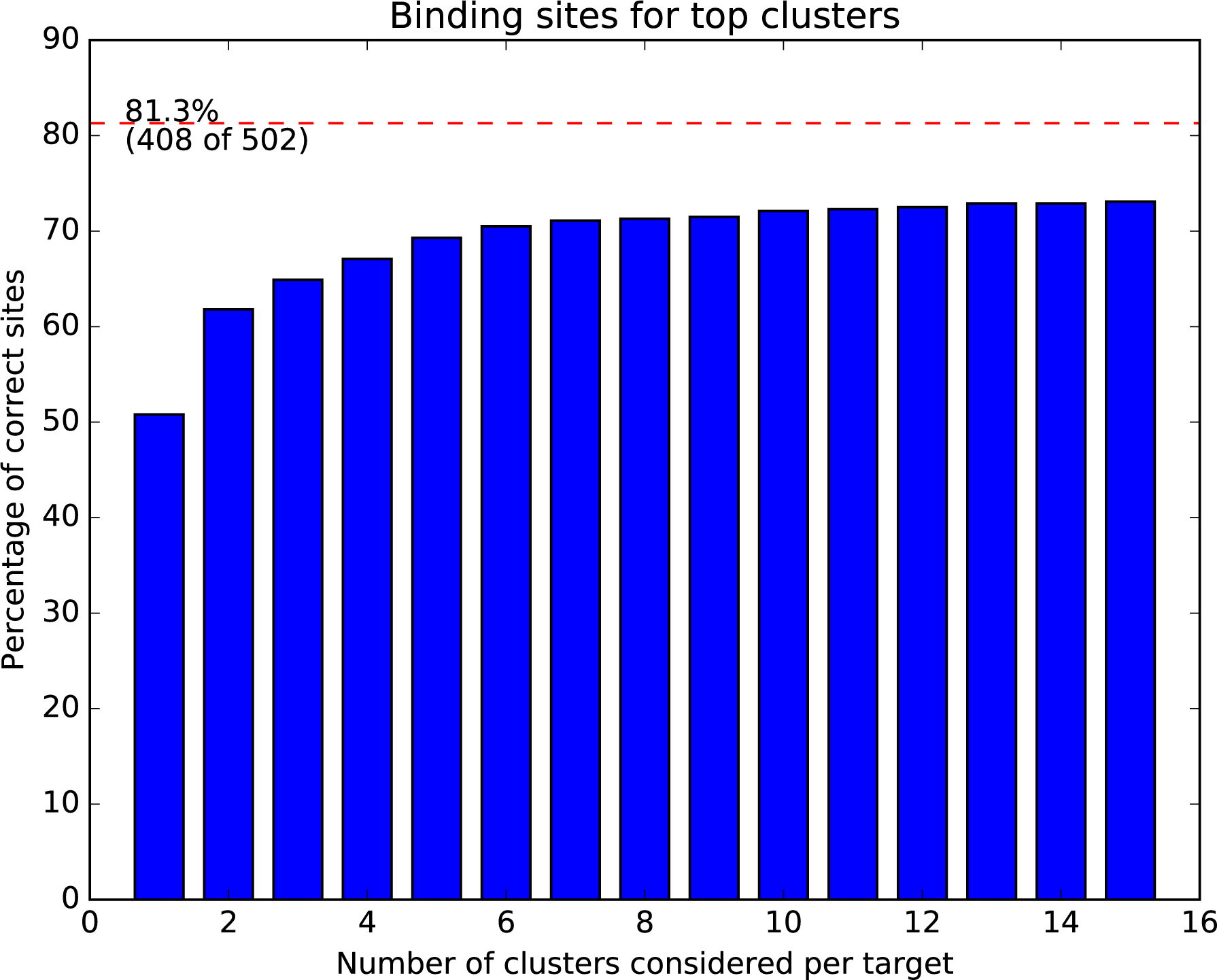
Fraction of correctly predicted binding sites when using the top ranked clusters. The red dotted line marks how many of the sites that are possible to describe using templates from the PDB, as previously shown in Table 2

## Conclusions

Overall, InterPep has proven a powerful tool for protein-peptide interaction site prediction. We show that a majority (81.3%) of protein-peptide interactions in the test set have structural templates without significant sequence similarity, and that a large proportion (67.7%) of the protein-peptide interactions actually have templates from protein-protein interactions. The InterPep pipeline, designed to take advantage of this, successfully identified half (50.7%) of the binding sites at top-1 and a majority (69.3%) among the top-5 using only structures with low sequence similarity as templates. In addition, the confidence score from InterPep correlates fairly well (0.63) with the precision of the prediction as measured by MCC for the predictions. By training InterPep also on modelled structures the performance is maintained for cases when the native structure is missing. In fact, the quality of the model did not turn out to be a determining factor for performance. InterPep is available as open source, and should be a useful tool in protein-peptide interaction analysis, providing possible guidelines for initial experiments or analysis of results. It should also be possible to use the results from InterPep as restraints to build 3D models of protein-peptide interaction.

## Acknowledgments

This work was supported by a Swedish Research Council grant, 2016-05369, The Swedish e-Science Research Center, and the Foundation Blanceflor Boncompagni Ludovisi, née Bildt. The computations were performed on resources provided by the Swedish National Infrastructure for Computing (SNIC) at the National Supercomputer Centre (NSC) in Linköping.

## Contributions

I.J. and B.W. designed the study. I.J. performed the method development. I.J. and BW performed data analysis. I.J. and B.W. wrote the manuscript with contributions from C.M. All authors reviewed the manuscript

## References

1. Mohan, A. et al. Analysis of molecular recognition features (MoRFs). J. Mol. Biol. 362, 1043–1059 (2006).

2. Diella, F. et al. Understanding eukaryotic linear motifs and their role in cell signaling and regulation. Front. Biosci. : A J. Virtual Libr. 13, 6580–6603 (2008).

3. Uversky, V. N. Intrinsically disordered proteins from A to Z. The Int. J. Biochem. & Cell Biol. 43, 1090–1103 (2011).

4. Uljon, S. et al. Structural basis for substrate selectivity of the E3 ligase COP1. Structure 24, 687–696 (2016).

5. Fischle, W. et al. Molecular basis for the discrimination of repressive methyl-lysine marks in histone H3 by Polycomb and HP1 chromodomains. Genes & development 17, 1870–1881 (2003).

6. Midic, U., Oldfield, C. J., Dunker, A. K., Obradovic, Z. & Uversky, V. N. Protein disorder in the human diseasome: unfoldomics of human genetic diseases. BMC Genomics 10 Suppl 1, S12 (2009).

7. Tu, W. B. et al. Myc and its interactors take shape. Biochimica et Biophys. Acta 1849, 469–483 (2015).

8. Vassilev, L. T. et al. In vivo activation of the p53 pathway by small-molecule antagonists of MDM2. Science 303, 844–848 (2004).

9. Hammoudeh, D. I., Follis, A. V., Prochownik, E. V. & Metallo, S. J. Multiple independent binding sites for small-molecule inhibitors on the oncoprotein c-Myc. J. Am. Chem. Soc. 131, 7390–7401 (2009).

10. Metallo, S. J. Intrinsically disordered proteins are potential drug targets. Curr. Opin. Chem. Biol. 14, 481–488 (2010).

11. Rhodes, G. *Crystallography made Crystal Clear*: A Guide for Users of Macromolecular Models (Academic press, 2010).

12. Puig, O et al. The tandem affinity purification (TAP) method: a general procedure of protein complex purification. Methods (San Diego, Calif.) 24, 218–229 (2001).

13. Parrish, J. R., Gulyas, K.D. & Jr, R.L.F. Yeast two-hybrid contributions to interactome mapping. Curr. Opin. Biotechnol. 17, 387–393 (2006). Protein technologies.

14. Roux, K. J., Kim, D. I. & Burke, B. BioID: a screen for protein-protein interactions. Curr. Protoc. Protein Sci. 19–23 (2013).

15. Shoemaker, B.A. & Panchenko, A. R. Deciphering protein-protein interactions. Part II. Computational methods to predict protein and domain interaction partners. PLoS Comput. Biol. 3, e43 (2007).

16. You, Z.-H., Lei, Y.-K., Zhu, L., Xia, J. & Wang, B. Prediction of protein-protein interactions from amino acid sequences with ensemble extreme learning machines and principal component analysis. BMC Bioinforma. 14 Suppl 8, S10 (2013).

17. Zhang, Q. C., Petrey, D., Garzón, J. I., Deng, L. & Honig, B. PrePPI: a structure-informed database of protein-protein interactions. Nucleic Acids Res. 41, D828–33 (2013).

18. Chen, H. & Skolnick, J. M-tasser: an algorithm for protein quaternary structure prediction. Biophys. J. 94, 918–928 (2008).

19. Mukherjee, S. & Zhang, Y. Protein-protein complex structure predictions by multimeric threading and template recombination. Structure 19, 955–966 (2011).

20. Sinha, R., Kundrotas, P.J. & Vakser, I. A. Docking by structural similarity at protein-protein interfaces. Proteins: Structure, Function, and Bioinformatics 78, 3235–3241 (2010).

21. Wallner, B. & Mirabello, C. Interpred: A pipeline to identify and model protein-protein interactions. Proteins: Structure, Function, and Bioinformatics 85, p1159–1170 (2017). DOI 10.1002/prot.25280.

22. Lensink, M. F. et al. The challenge of modeling protein assemblies: The CASP12-CAPRI experiment. Proteins: Structure, Function, and Bioinformatics 86, 257–273 (2017).

23. Berman, H. M. et al. The protein data bank. Nucleic Acids Res. 28, 235–242 (2000).

24. Hetényi, C. & van der Spoel, D. Efficient docking of peptides to proteins without prior knowledge of the binding site. Protein Sci. 11, 1729–1737 (2002).

25. Raveh, B., London, N. & Schueler-Furman, O. Sub-angstrom modeling of complexes between flexible peptides and globular proteins. Proteins: Structure, Function, and Bioinformatics 78, 2029–2040 (2010).

26. Raveh, B., London, N., Zimmerman, L. & Schueler-Furman, O. Rosetta flexpepdock ab-initio: simultaneous folding, docking and refinement of peptides onto their receptors. PloS One 6, e18934 (2011).

27. Kurcinski, M., Jamroz, M., Blaszczyk, M., Kolinski, A.& Kmiecik, S. CABS-dock web server for the flexible docking of peptides to proteins without prior knowledge of the binding site. Nucleic Acids Res. 43, W419–24 (2015).

28. Petsalaki, E., Stark, A., García-Urdiales, E. & Russell, R. B. Accurate prediction of peptide binding sites on protein surfaces. PLoS Comput. Biol. 5, e1000335 (2009).

29. Trabuco, L. G., Lise, S., Petsalaki, E. & Russell, R. B. Pepsite: prediction of peptide-binding sites from protein surfaces. Nucleic Acids Res. 40, W423–W427 (2012).

30. Lavi, A. et al. Detection of peptide-binding sites on protein surfaces: the first step toward the modeling and targeting of peptide-mediated interactions. Proteins 81, 2096–2105 (2013).

31. Saladin, A. et al. PEP-SiteFinder: a tool for the blind identification of peptide binding sites on protein surfaces. Nucleic acids research 42, W221–6 (2014).

32. Yan, C. & Zou, X. Predicting peptide binding sites on protein surfaces by clustering chemical interactions. Journal of Computational Chemistry 36, 49–61 (2015).

33. Brenke, R. et al. Fragment-based identification of druggable ‘hot spots’ of proteins using Fourier domain correlation techniques. Bioinforma. (Oxford, England) 25, 621–627 (2009).

34. Lee, H., Heo, L., Lee, M. S. & Seok, C. GalaxyPepDock: a protein-peptide docking tool based on interaction similarity and energy optimization. Nucleic acids research 43, W431–5 (2015).

35. Alam, N. et al. High-resolution global peptide-protein docking using fragments-based PIPER-FlexPepDock. PLoS computational biology 13, e1005905 (2017).

36. Kozakov, D., Brenke, R., Comeau, S. R. & Vajda, S. PIPER: an FFT-based protein docking program with pairwise potentials. Proteins 65, 392–406 (2006).

37. Vanhee, P. et al. Protein-peptide interactions adopt the same structural motifs as monomeric protein folds. Structure 17, 1128–1136 (2009).

38. Gao, M. & Skolnick, J. Structural space of protein-protein interfaces is degenerate, close to complete, and highly connected. PNAS 107, 22517–22522 (2010).

39. Kundrotas, P. J., Zhu, Z., Janin, J. & Vakser, I. A. Templates are available to model nearly all complexes of structurally characterized proteins. Proc. Natl. Acad. Sci. 109, 9438–9441 (2012).

40. Hubbard, S.J. & Thornton, J. M. Naccess. Computer Program, Department of Biochemistry and Molecular Biology, University College London2 (1993).

41. Alva, V., Nam, S.-Z., Söding, J. & Lupas, A. N. The mpi bioinformatics toolkit as an integrative platform for advanced protein sequence and structure analysis. Nucleic Acids Res. 44, W410–W415 (2016).

42. Alva, V., Nam, S.-Z., Söding, J. & Lupas, A. N. The mpi bioinformatics toolkit as an integrative platform for advanced protein sequence and structure analysis. Nucleic Acids Res. 44, W410–W415 (2016).

43. Webb, B. & Sali, A. Comparative Protein Structure Modeling Using MODELLER. Current Protocols in Protein Science 86, 2.9.1–2.9.37 (2016).

44. Zhang, Y. & Skolnick, J. TM-align: a protein structure alignment algorithm based on the TM-score. Nucleic acids research 33, 2302–2309 (2005).

45. Zhang, Y. & Skolnick, J. Scoring function for automated assessment of protein structure template quality. Proteins: Structure, Function, and Bioinformatics 57, 702–710 (2004).

46. Pedregosa, F. et al. Scikit-learn: Machine learning in Python. Journal of Machine Learning Research 12, 2825–2830 (2011).

47. Jones, E., Oliphant, T., Peterson, P. et al. SciPy: Open source scientific tools for Python (2001).

48. Ketchen Jr, D. J. & Shook, C. L. The application of cluster analysis in strategic management research: an analysis and critique. Strateg. management journal 441–458 (1996).

49. Mayrose, I., Graur, D., Ben-Tal, N. & Pupko, T. Comparison of site-specific rate-inference methods for protein sequences: empirical bayesian methods are superior. Mol. biology evolution 21, 1781–1791 (2004).

50. London, N., Movshovitz-Attias, D. & Schueler-Furman, O. The structural basis of peptide-protein binding strategies. Structure 18, 188–199 (2010).

51. Cristobal, S., Zemla, A., Fischer, D., Rychlewski, L. & Elofsson, A. A study of quality measures for protein threading models. BMC Bioinforma. 2, 5 (2001).

52. Illergård, K., Ardell, D.H. & Elofsson, A. Structure is three to ten times more conserved than sequence-a study of structural response in protein cores. Proteins: Structure, Function, and Bioinformatics 77, 499–508 (2009).

53. Petsko, G.A. & Ringe, D. Protein structure and function (New Science Press, 2004).

